# Nucleolar Targeting and ROS-dependent inhibition of rRNA synthesis by Epstein-Barr Virus Nuclear Antigen 1

**DOI:** 10.64898/2025.12.29.696898

**Authors:** M. Chabi, B. Aguida, T. Laudat, K. Villette, N. Oufella, S. Castro Da Costa, V. Stierlé, V. Sirri, P. Roussel, C. Akpovi, J. Pothier, N. Jourdan

## Abstract

Epstein–Barr virus nuclear antigen 1 (EBNA-1) is essential for viral episome maintenance and is consistently expressed in all forms of EBV latency. In a previous study, we observed EBNA-1 localizing in the nucleolus, as well as its interaction with the nucleolar protein EBP2 in both the nucleoplasm and nucleoli. Nucleoli are dynamic nuclear domains that coordinate ribosome assembly with the regulation of cell cycle progression and stress responses. Here, we studied the mechanism by which EBNA-1 achieves nucleolar localization and functional consequences for host cell physiology. We observed that EBNA-1 nucleolar accumulation occurs specifically during late G1 and S phase and depends on a bipartite nucleolar localization signal (NoLS) composed of two cooperative Weber motifs separated by 24 amino acids. Mutations disrupting NoLS abolish interaction with the nucleolar protein EBP2, whose depletion similarly prevents EBNA-1 nucleolar import. Once in the nucleolus, EBNA1 triggers a 50 % reduction in the synthesis of ribosomal RNA (rRNA) which leads to a global reduction in cellular protein production. In addition, EBNA1 nucleolar targeting induces reactive oxygen species (ROS) production and antioxidant treatment restores both rRNA transcription and protein synthesis. These findings identify EBP2 as a viral nucleolar docking partner and establish nucleolar targeting of EBNA1 as a regulated, cell cycle–dependent process that links oxidative stress to the perturbation of ribosome biogenesis. This mechanism may contribute to EBNA1’s ability to promote cell survival and transformation.

**IMPORTANCE:** EBNA1 is the only Epstein–Barr virus protein that is consistently expressed in all forms of latency and in every EBV-associated tumor. Although it localizes to the nucleolus and interacts with three nucleolar proteins, its influence on nucleolar functions had never been investigated. Here, we demonstrate that EBNA1’s nucleolar targeting is a strictly cell-cycle–regulated process that depends on a previously unrecognized bipartite nucleolar localization signal as well as its interaction with the nucleolar protein EBP2. Once inside the nucleolus, EBNA1 induces a ROS-dependent suppression of rRNA synthesis, resulting in a global decrease in cellular protein production. In addition, these findings uncover a new function of EBNA1: the ability to induce nucleolar stress without activating canonical apoptotic pathways, thereby promoting host-cell survival and potentially contributing to EBV-driven oncogenesis

## INTRODUCTION

Epstein-Barr virus (EBV), a gamma herpesvirus, infects over 95% of the global population. Like all members of the *Herpesviridae* family, it establishes a lifelong latent infection following primary infection. Primary infection is usually asymptomatic in infancy but can result in infectious mononucleosis, a self-limiting disease, when contracted during adolescence or adulthood. During latency, however, EBV can become highly oncogenic. Indeed, several studies have demonstrated the involvement of EBV in various malignant tumors, such as Burkitt’s lymphoma, Hodgkin’s disease, nasopharyngeal carcinoma, certain gastric adenocarcinomas (1), as well as posttransplant lymphoproliferative disorder (2).

The virus is transmitted through saliva and crosses the oropharyngeal epithelial barrier to infect naïve B cells in the lymphoid tissue. During this infection, the viral genome, a 171,820 bp linear DNA molecule, forms a circular episome and sequentially expresses three latency gene programs. These drive B cell activation (latency III) and maturation to the memory stage (latency II) and lifelong persistence (latency I) in memory B cells within the bloodstream (3, 4).

EBV Nuclear Antigen 1 (EBNA-1) is a nuclear protein expressed across all three latency programs and is the only viral protein present in latency I. EBNA-1 plays a crucial role in viral persistence, mediating transcription, replication and tethering of viral episome to host chromatin throughout the cell cycle of dividing B lymphocytes (5). In addition to its functions during the latent viral cycle, EBNA-1 was reported to induce tumors in transgenic mice (6) and enhance B cell line immortalization by several thousand fold (7). Consistently, EBNA-1 knockdown reduces the proliferation and survival of EBV-positive tumor cell lines (8–11). Although the mechanisms are not yet fully understood, EBNA-1 may promote oncogenesis by inhibiting p53-dependent apoptosis (8, 12) and upregulating surviving expression (13).

Previous studies have observed the localization of EBNA-1 in the nucleoli of transfected cells (14, 15), as well as its interaction with the nucleolar protein EBP2 (16) in both the nucleoplasm and nucleoli (15). Nucleoli are the major nuclear compartments of the cell nucleus where ribosomal RNAs (rRNAs) are synthesized, processed and assembled into ribosome subunits. Nucleoli can also act as stress cellular sensors (17) by detecting DNA damage, thermal shock or ribosome synthesis defects. Such perturbations trigger the so-called nucleolar stress that start by an inhibition of rRNA transcription, associated with the delocalization of nucleolar proteins to the nucleoplasm. Delocalized nucleolar proteins of which B23 (also known as nucleophosmin or NMP1) is the main one, bind to the P53 inhibitor MDM2, leading to activation of the tumor suppressor p53 whose activity will cause cell cycle arrest or apoptosis (18).

In the last few years, a growing list of studies has shown that viruses, regardless of the nature of their genome or the cytoplasmic or nuclear localization of their replication cycle, produce proteins that accumulate in the nucleolus during infection or following transfection, thereby altering nucleolar functions, and influencing the viral replication cycle and pathogenesis (for a review see (19)). There are numerous examples, notably in *Herpesviridae* family, where viral proteins can subvert or inactivate nucleolus components to ensure their lytic replication (20–28).

Although three nucleolar proteins, nucleolin (29), B23 (30, 31) and EBP2 (16), have already been identified as EBNA-1 or EBNA-1 mRNA interaction partners, the involvement of nucleoli in EBV viral cycle and pathophysiology remains unexplored. Here we sought to investigate the mechanism behind EBNA-1’s nucleolar localization and its potential impact on nucleolar functions.

## RESULTS

### Heterogenous nucleolar staining

The nucleolus is a membrane-less organelle organized in a tripartite structure that consist of several fibrillar center (FC), each surrounded by a dense fibrillar component (DFC), the whole surrounded by the granular component (GC). In order to study EBNA-1 nucleolar localization carefully, a large number of HeLa cells co-transfected with EBNA-1-GFP and DsRed-tagged EBP2, were observed by live cell imaging during interphase. Nucleolar EBNA-1 staining appeared to be heterogeneous from cell to cell. In 80% of the cells, the distribution of GFP-EBNA-1 covered the entire area of the nucleolus, colocalized with almost 100% of the EBP2 staining pattern (Fig. 1, panels 1-3) that corresponds to the GC pattern.

**Fig. 1.**
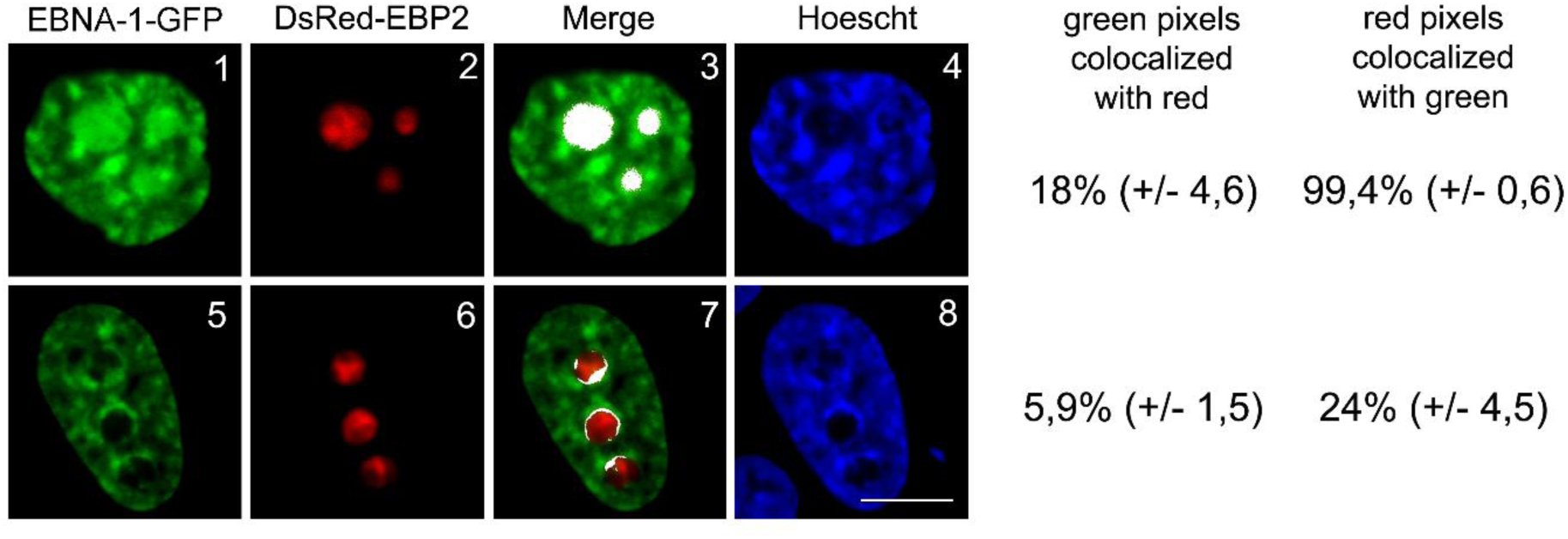
Nuclear and nucleolar localization of EBNA-1-GFP by confocal microscopy in living cells. HeLa cells were co-transfected with EBNA-1-GFP and DsRed–EBP2. The DNA was stained with Hoechst. Images 1 to 8 show single confocal z-section. Colocalized pixels (in white) from the merge images were quantified with ImageJ software. The standard deviations are indicated (more than 200 cells were analyzed across 12 independent experiments). Scale bar, 10 μm.

Conversely, in 20% of the cells, GFP-EBNA-1 exhibited a slightly different nucleolar pattern, where it concentrated in areas surrounding nucleoli and partially overlapping with the periphery of the region occupied by EBP2 (Fig. 1, panels 5-7). This perinucleolar pattern of EBNA-1 did not correspond to any nucleolar compartment but rather perfectly matches with perinucleolar heterochromatin as observed by Hoechst staining (Fig. 1, panel 8). Indeed nucleoli were frequently surrounded by a layer of heterochromatin, thus EBNA-1 perinucleolar staining is consistent with chromatin binding property of EBNA-1 conferred by three independent binding sites (CBS1, 2 and 3) (32). Moreover, the fact that perinucleolar heterochromatin makes interdigitations toward the interior of nucleolus (33, 34) might well explain EBNA-1 colocalization with the periphery of the EBP2 staining pattern. These two different EBNA-1 staining patterns will be referred as to nucleolar and peri-nucleolar patterns thereafter.

### Cell cycle regulation of EBNA-1 nucleolar targeting

Heterogeneity observed for EBNA-1 nucleolar staining in asynchronous cells could suggest a cell-cycle regulation of EBNA-1 nucleolar targeting. To test this hypothesis, HeLa cells co-transfected with EBNA-1-GFP and DsRed-B23, a nucleolar protein also known as NPM or nucleophosmin, were synchronized as described in materials and methods section and observed at different phase of the cell cycle. The pre-rRNA processing factor B23 labels nucleoli from the appearance of prenucleolar bodies (PNBs) in telophase (Fig. 2, panel b), which progressively dissipate in early G1 as they are incorporated into the reforming nucleoli, and continues to mark them throughout late G1, S and G2 (Fig. 2, panels e, h and k), until their disassembly at mitotic onset.

**Fig. 2.**
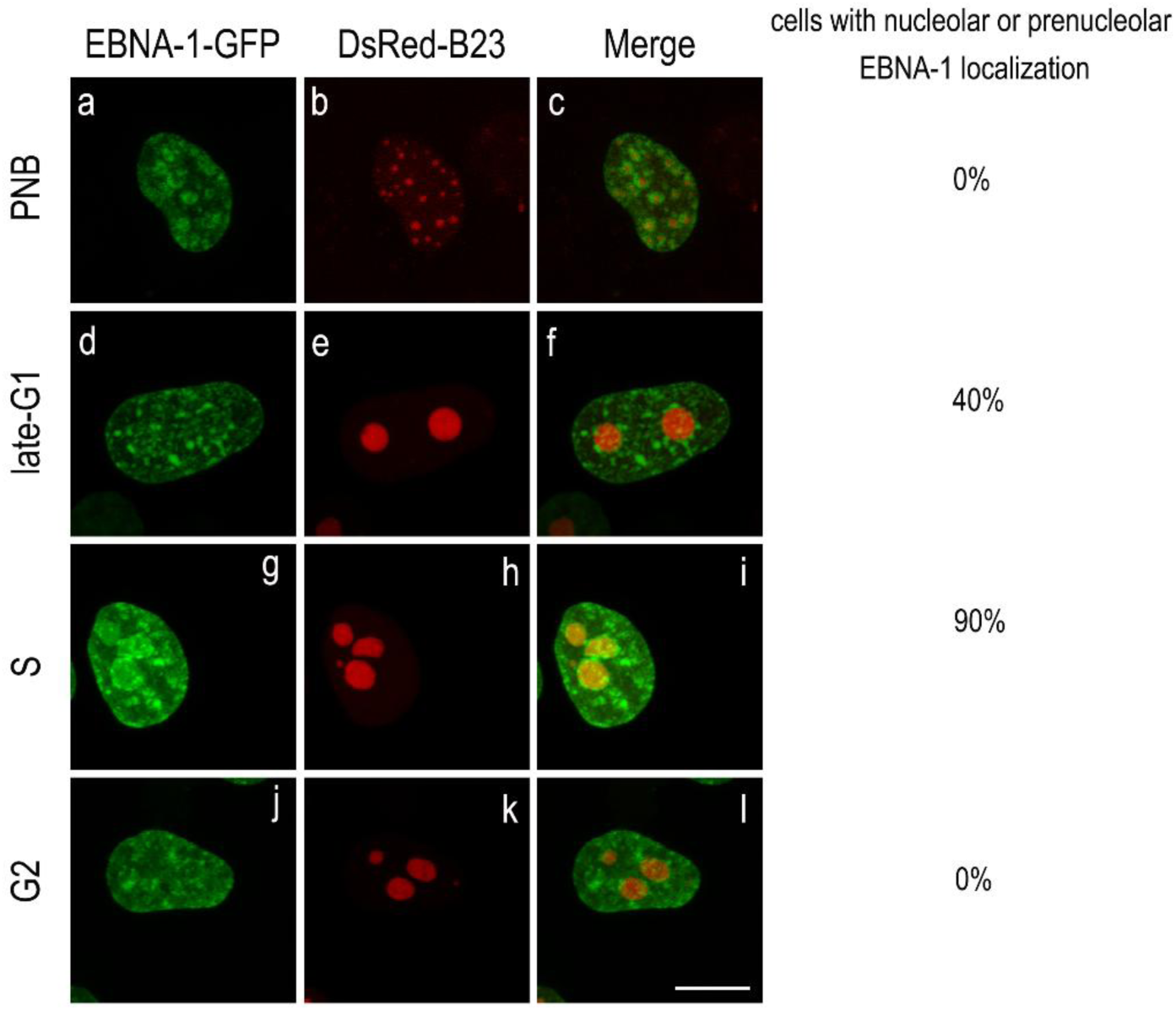
Subcellular localization of EBNA-1-GFP at various stages of the cell cycle. HeLa cells were co-transfected with EBNA-1-GFP and DsRed-B23 and subjected either to cell-cycle synchronization combined with RNA polymerase I inhibition to visualize PNBs, or to a double-thymidine block followed by release to obtain cells in late G1, S, and later G2 phases. Images a to l show single confocal z-section and are representatives of an average of 50 cells observed at each stage across 3 independent experiments. Scale bars, 10 μm.

Although EBNA-1 is detected throughout the cell cycle on heterochromatin surrounding prenucleolar bodies (Fig. 2, panels a and c) and nucleoli (panels d, f, g, i, j, and l), no EBNA-1 signal is observed within prenucleolar bodies (PNBs) or within nucleoli during G2. Instead, nucleolar localization of EBNA-1 becomes evident in late G1 in approximately 60% of transfected cells and reaches about 90% during S phase (Fig. 2, panels g–i). These findings indicate that EBNA-1 nucleolar targeting is tightly cell-cycle regulated and peaks during the DNA synthesis phase.

### The EBP2 binding site is a GAR related domain

Interaction with nucleolar proteins, rRNAs or rDNAs had been proposed as a mechanism for nucleolar targeting of proteins (35, 36). According to that hypothesis, a nucleolar localization signal (NoLS) would be a nucleolar molecule interacting sequence. Considering that direct binding of EBNA-1 to EBP2 have been well characterized (14, 16) and moreover have been localized in nucleoplasm as well as in nucleoli (15), it may be postulated that the latter transports EBNA-1 to the nucleoli. Taking advantage of the nucleolar database enhancement (37) due to organelle-directed mass spectroscopy studies (38), we analysed amino acids computational feature of the 325 to 376 amino acids sequence defined as the EBP2 binding region by Wu and colleagues (14). Interestingly, the EBP2 binding region (or CBS2, see Fig. 3A) contains Gly-Arg rich motifs, most particularly a GRGRGG motif that is repeated three times in a short region. This observation led us to compute occurrences of all tripeptides and hexapeptides containing mixed Arg and Gly residues in the human nucleolar and non-nucleolar proteome (as explained in the “Materials and Methods” section). As described in Fig. 3B, the tripeptides GRG and RGG were enriched in the human nucleolar proteome 1.43 and 1.5-fold respectively as compared with the non-nucleolar human proteome, and nearly 25% of the human nucleolar proteins contained one of these tripeptides at least once. Moreover, the ratio of nucleolar proteins containing the hexapeptide GRGRGG is about 7 times that of non-nucleolar proteins and, similarly to the EBP2 binding site of EBNA-1, fibrillarin, one of the major nucleolar proteins, contains this hexapeptide in closely repeated three copies. Interestingly, the three repeated hexapeptide motifs are included in a longer GR rich region, which matches exactly the EBP2 binding site (Fig. 3C).

**Fig 3.**
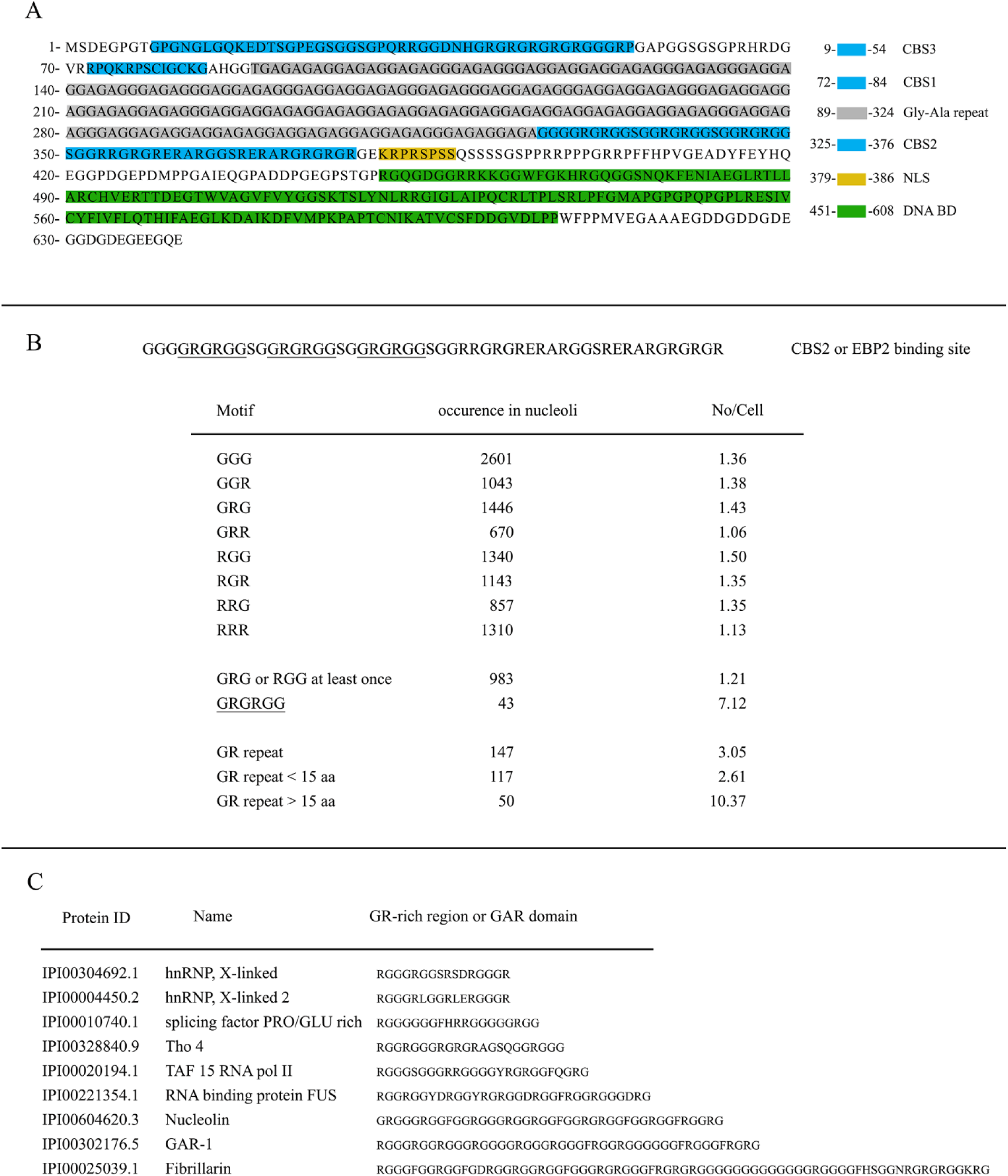
EBNA-1 sequence analysis. (A) **Functional mapping of EBNA-1** (B) occurrence of different combinations of short or long GR repeats in the Nucleolar Human proteome database (“occurrence in nucleoli”) and relative abundance in the Nucleolar Human proteome versus in the Human proteome database (“No/Cell”). (C) Nucleolar proteins with motifs at least 7 amino-acids long and 80% of GR repeat composed at least of 20% G or R.

Then, we designed an algorithm (see “Materials and Methods” section) that allowed us to search for such GR rich regions in nucleolar and non-nucleolar proteomes. GR rich regions were enriched 3-fold in the human nucleolar proteome as compared with non-nucleolar human proteome, moreover enrichment increased to 10-fold for GR rich region longer than 15 amino acids (Fig. 3B).

Examples of nucleolar proteins containing such a long GR rich region are listed in Fig. 3C and included nucleolin, fibrillarin, Tho4, and several heterogeneous nucleolar ribonucleoproteins (hnRNP) all involved either in rRNA processing or in mRNA transcription, processing and export. These long GR-rich domains have been defined, at least for nucleolin and fibrillarin (39) and GAR-1 (40) as GAR (for glycine arginine rich) domain.

### Two Weber’s motif constitute the EBNA-1 NoLS

GAR domains have been reported to interact with ribosomal proteins (41), RNA or DNA (42) and may function as nucleolar localization signals (NoLS) (43). To assess their contribution to the nucleolar targeting of EBNA-1, we generated an EBNA-1 mutant lacking the entire GAR domain (Δ325–376) fused to GFP, and compared its nucleolar localization with that of the GFP-EBNA-1 fusion protein. As shown in Fig. 4 (panels a–f), deletion of the GAR domain reduced by 75% the proportion of cells displaying EBNA-1 nucleolar localization. These findings indicate that the GAR domain is critical for the nucleolar localization of EBNA-1 and strongly support its role as a NoLS.

**Fig. 4.**
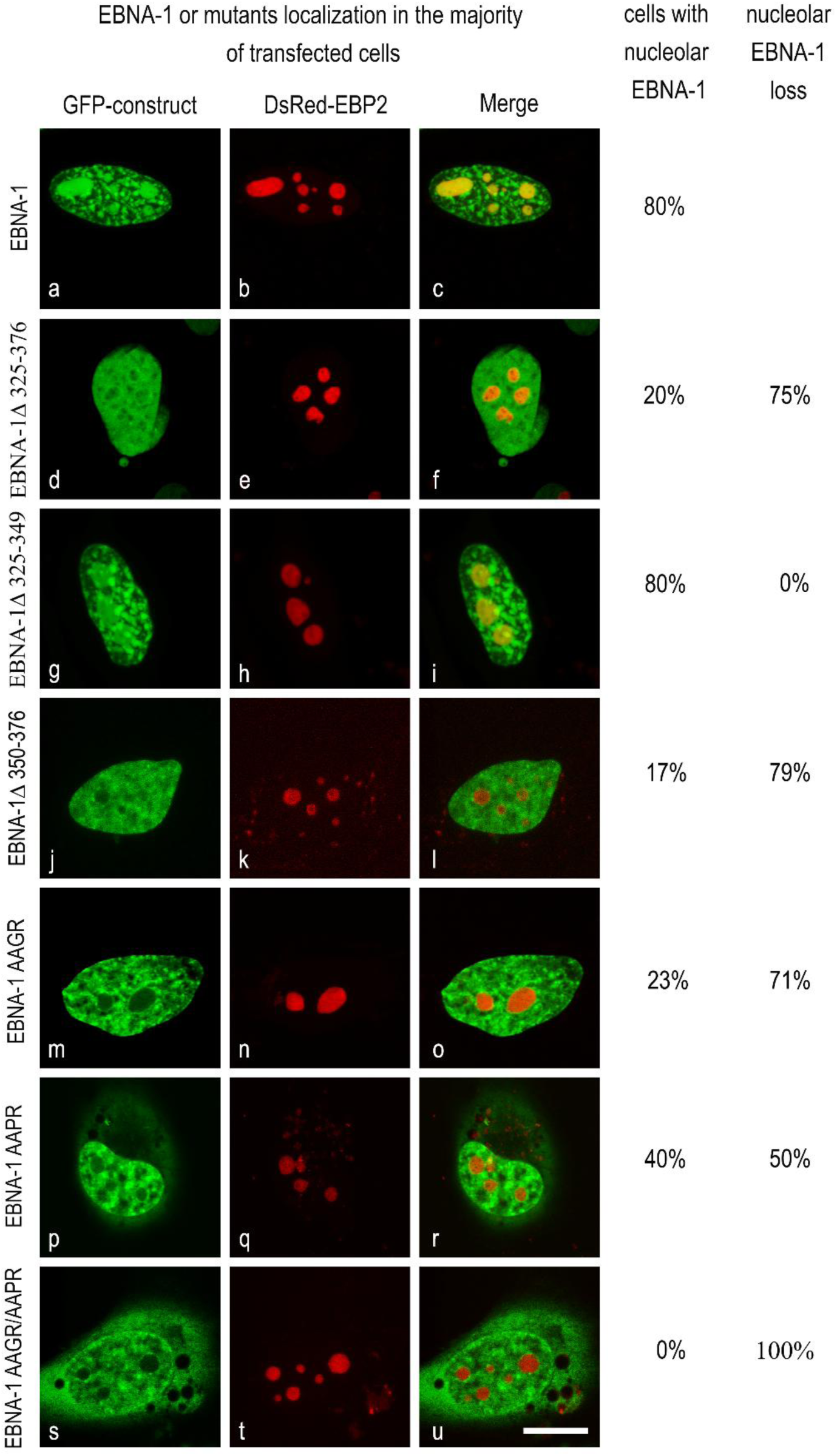
Localization of EBNA-1 and mutants in transfected cells. HeLa cells were co-transfected with EBNA-1-GFP or EBNA-1-mutant-GFP and DsRed–EBP2. Images a to u show single confocal z-section. Scale bar, 10 μm. Percentages were calculated from >100 observations for each mutant across 12 independent experiments.

To further delineate the NoLS within the GAR domain, we generated EBNA-1 mutants lacking the three tandem GRGRGG repeats (Δ325–349). As shown in Fig. 4 (panels g-i), this deletion did not impair EBNA-1 nucleolar localization, as the protein still colocalized with EBP2 in 80% of analyzed cells. These findings indicate that, although the GAR domain contributes to nucleolar targeting, this property cannot be attributed to the GRGRGG repeats. In contrast, deletion of the remaining portion of the GAR domain (Δ350–376) led to a 79% reduction in nucleolar localization (Fig. 4, panels j-l).

Interestingly, the 350-376 part of the GAR domain possesses a K/R-K/R-X-K/R Weber’s motif, the RRGR sequence at position 351 separated by 24 aa from a second Weber’s motif KRPR at position 379 in the NLS (see Fig. 3A). Although no canonical NoLS has been defined, some are arranged around Weber’s motif in one or multiple copies (44). In order to clearly identify the EBNA-1 NoLS, we performed site directed mutagenesis at the 351 or at the 379 or at both Weber’s motifs replacing the first two basic aa of each Weber’s motif to alanine as indicated in Fig. 4A. Whereas single mutants AAGR or AAPR are still present in nucleoli of respectively, 23% and 40 % of HeLa transfected cells, the nucleolar localization of the double mutant is totally abolished for 100 % of transfected Hela cells (Fig. 4, panels s to u). These results indicate that both motifs are necessary and work cooperatively to translocate EBNA-1 to nucleoli. Therefore, the NoLS of EBNA-1, described for the first time, is a bi-partite signal composed of two Weber’s motifs separated by 24 aa. It is noteworthy that this mutant, lacking the NoLS and hereafter referred to as EBNA1-NoLS*, specifically lost its nucleolar localization while retaining both nucleoplasmic and perinucleolar chromatin binding. This feature will later allow us to use this mutant as a reference control to investigate the effect of EBNA1 entry into the nucleolus.

### Mutation of the NoLS abolishes EBNA-1/EBP2 interaction

The EBNA-1 NoLS partially overlaps, but does not fully coincide, with the amino acid sequence 325–376 previously defined as the EBP2-binding region (14). Therefore, the hypothesis proposed above, that EBP2 acts as the docking partner mediating EBNA-1 transport to the nucleolus, appears less straightforward and requires experimental validation. For that we used Förster resonance energy transfer (FRET). FRET is a nonradiative energy transfer that can occur when a donor and a compatible acceptor fluorophore are located at a distance lower than 10 nm from each other (45). Due to this property, FRET can be used to monitor spatiotemporal protein-protein interaction in living cells. In a previous work, by using GFP-EBP2 as a donor and DsRed-EBNA-1 as an acceptor, we demonstrated that EBNA-1 interact with EBP2 in nucleoplasm and in nucleoli in living cells (15). Thus, to assess whether EBP2 serves as the docking partner of EBNA-1 for nucleolar translocation, we still used the GFP-EBP2/DsRed-EBNA-1 pair as a FRET-positive control. We then examined whether mutations that abolish the NoLS of EBNA-1 affect ability of the mutant (referred as EBNA-1-NoLS*) to interact with EBP2. In the present case, after excitation of the donor at 488 nm by an argon laser, FRET has been detected by analysis of donor and acceptor emission spectra between 498 and 648 nm. Fig. 5 C shows emission spectrum of the positive control with a double peaks shape at 513 nm and 588 nm that respectively correspond to GFP and DsRed emission. To confirm that peak at 588 nm results only from DsRed emission and that emission is due to DsRed excitation by FRET and not by a direct DsRed excitation by the argon laser, we performed two negative controls. The first one consists in the use of cells expressing the donor, GFP-EBP2, alone. After excitation at 488 nm, emission spectrum analysis (Fig. 5 A), show a typical GFP peak at 513 nm and no GFP emission at 588 nm, clearly indicating there is no GFP emission contribution in the DsRed detection channel at 588 nm. The second one consists in the use of cells expressing the acceptor, DsRed-EBNA-1, alone. As shown in the Fig. 5B, 488 nm excitation induce a very low intensity DsRed emission at 588 nm. This emission resulting from a direct DsRed excitation is hardly detected (or negligeable) with a mean fluorescence intensity of 12.33 +/- 3, compared with emission resulting from FRET in the positive control that present a mean fluorescence intensity of 167.36 +/- 36 (Fig. 5C). These two negative controls confirmed the occurrence of FRET between GFP-EBP2 and DsRed-EBNA-1 in living cells, which is consistent with our previously published results (15). To semi-quantitatively evaluate emission spectral changes, from different levels of protein expression in different cells, the mean ratio of fluorescence intensity (FI 588 nm/ FI 588 nm + FI 513 nm) detected from each nucleus has been determined and was 0.4 +/- 0.1 as shown in Fig. 5F. Thus, the double peaks spectral pattern and mean ratio of fluorescence intensity of 0.4 can be used as a positive FRET sign in the following analyses. Fig. 5D shows the GFP-EBP2/ EBNA-1-NoLS*-DsRed spectral patterns, obtained from 13 entire nuclei. In strong contrast to the positive control, they did not present prominent peak at 588 nm (Fig. 5D), and the mean ratio of fluorescence intensity decreased significantly to a level of 0.13 +/- 0.06, clearly indicating that FRET did not occur.

**Fig. 5.**
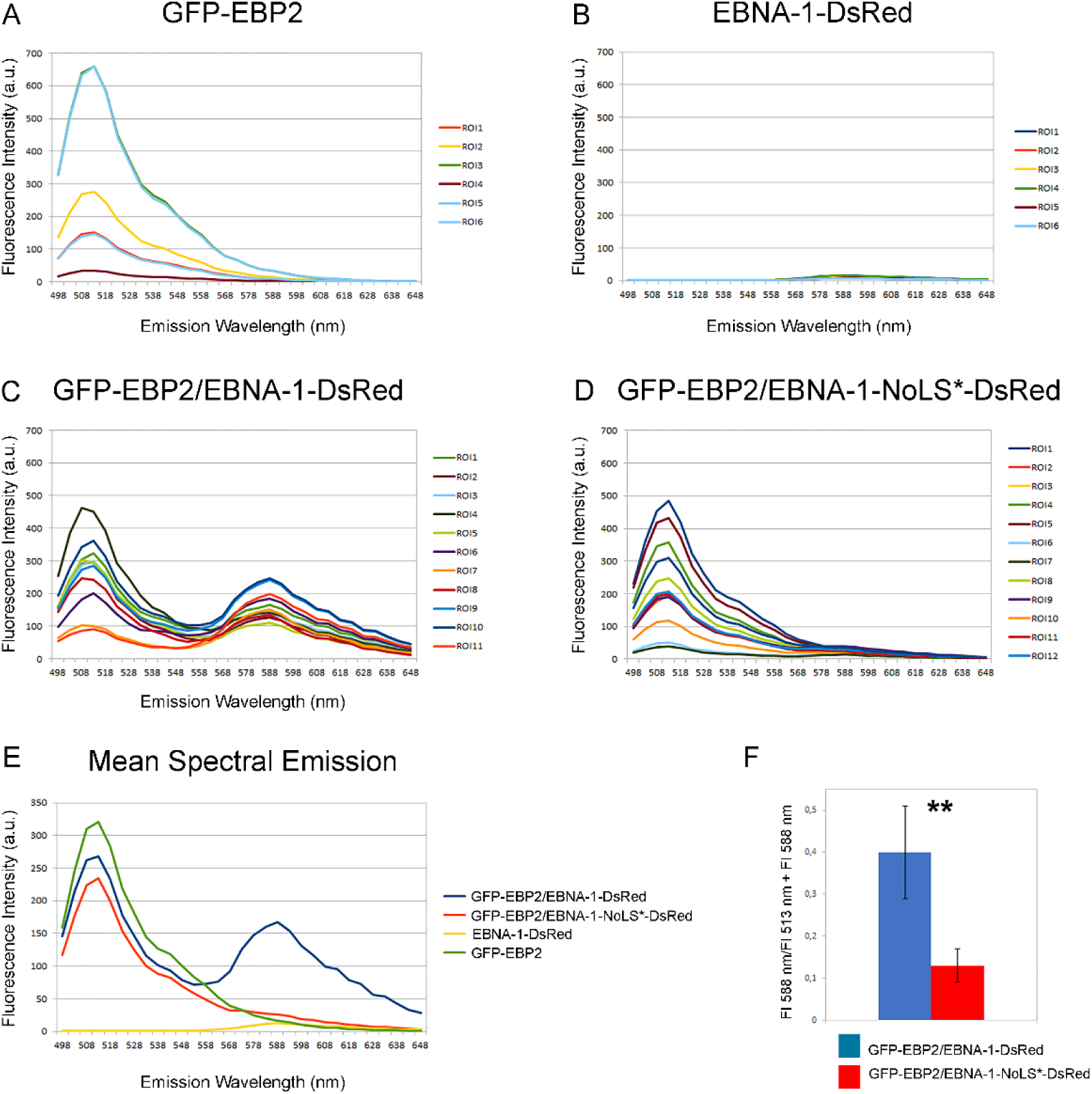
NoLS mutation abolish EBNA-1/ EBP2 interaction detected by FRET in living cells through spectral analysis. HeLa cells transfected with (A) GFP-EBP2, (B) EBNA-1-DsRed, or co-transfected with (C) GFP-EBP2 and EBNA1-DsRed, (D) GFP-EBP2 and EBNA1-NoLS*-DsRed were analyzed using an inverted TCS-SP5 confocal microscope. Fluorescence Intensity (FI) emissions between 498 and 648 nm were recorded after excitation at 488 nm in regions of interest (ROIs) corresponding to the nuclei of transfected or co-transfected cells. Data were processed using Calc (LibreOffice). (E) Mean fluorescence emission spectra from figures A to D. (F) Mean intensity ratios (R = FI 588 nm / [FI 513 nm + FI 588 nm]) calculated from the 513 nm and 588 nm peaks of each emission spectrum shown in figures C and D. Data are shown as mean *± SE.* The asterisks indicate significance level of the differences: **p-value < 0.01.

All these results indicate that mutations that abolish the NoLS also abolish interaction with EBP2, strongly suggesting that NoLS might mediate EBNA-1 interaction to EBP2 and that EBP2 might transport EBNA-1 to the nucleoli.

### siRNA-mediated knockdown of EBP2 reduced EBNA-1 nucleolar localization

To further substantiate the role of EBP2 in EBNA-1 nucleolar import, EBP2 expression in HeLa cells was silenced using siRNA. Knockdown efficiency was assessed 24 h and 48 h post-siRNA transfection by RT-qPCR and Western blotting. RT-qPCR analysis revealed a sustained reduction in EBP2 mRNA detectable at 24 h and maintained through 48 h (Fig. 6A). Consistently, Western blot analysis confirmed these results, showing a parallel decrease in EBP2 protein levels with a maximum efficiency at 48h post siRNA (Fig. 6B).

**Fig. 6.**
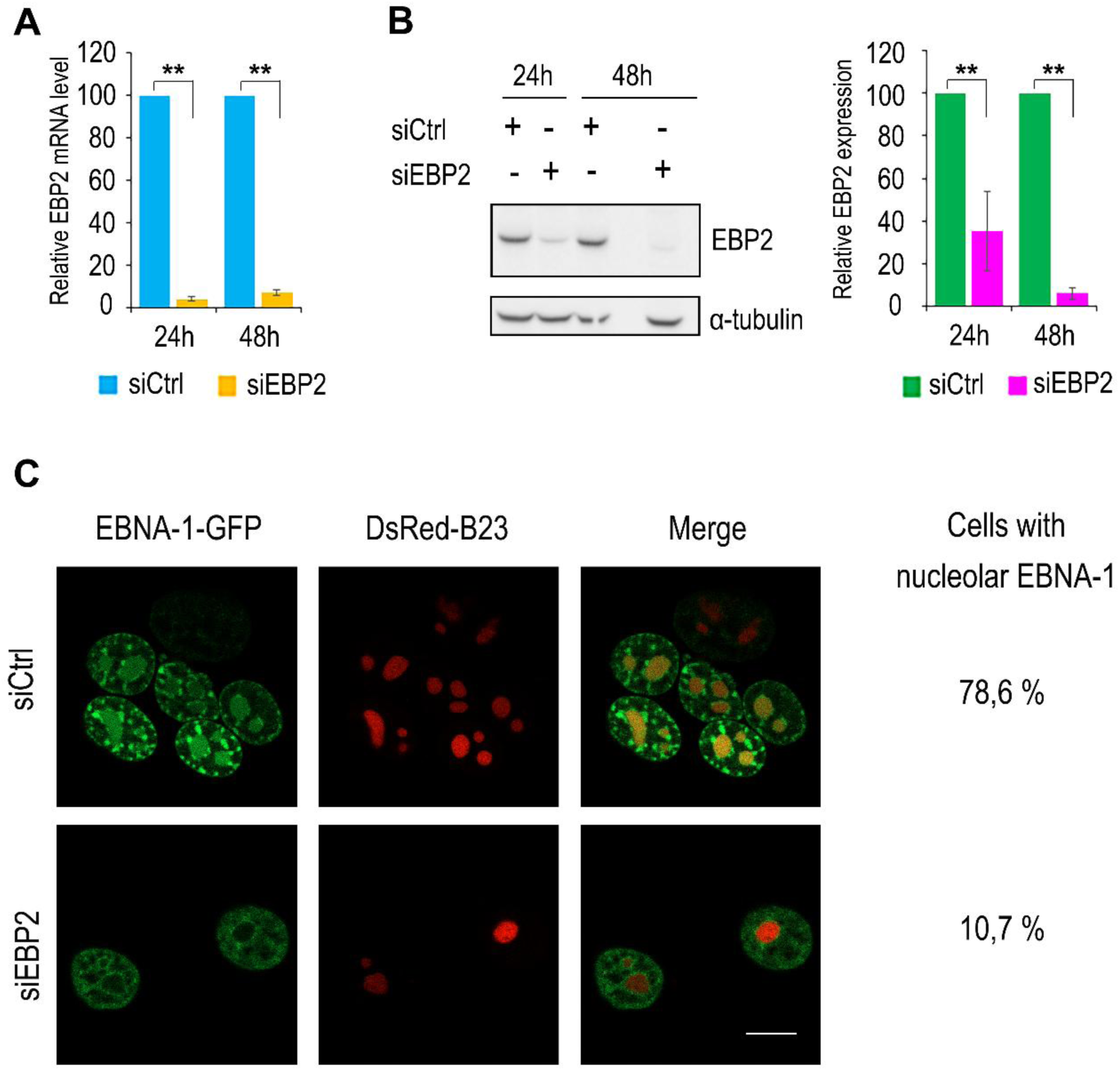
EBP2 silencing using siRNA effectively suppresses EBNA-1 nucleolar targeting. (A and B) HeLa cells transfected with siRNA control (siCtrl) or the EBP2-targeting siRNA (siEBP2) were harvested at 24h and 48h post transfection and analyzed for EBP2 silencing efficacy (A) Analysis at mRNA expression levels detected by RT-qPCR. Significance thresholds were determined by one-way ANOVA comparing test conditions with the control group, using biological and technical triplicates (**p < 0.001) (B) Analysis at protein expression levels by Western blot. Densitometric analyses from three independent experiments are shown as histograms. Significance thresholds were determined by one-way ANOVA comparing test conditions with the control group (**p < 0.001). (C) 24h post transfection with siCtrl or with 4 distinct siEBP2, Hela cells were co-transfected with EBNA-1-GFP and DsRed-B23 and observed by confocal microscopy 48h post-siRNA transfection. Images show single confocal z-section. Scale bar, 10 μm. Percentages were calculated from >100 observations for each condition across 3 independents experiments.

Since our objective was to assess how EBP2 silencing affects EBNA-1 targeting to nucleoli, EBNA-1-GFP was transfected at 24 h, and its localization was examined at 48 h, when EBP2 expression reached its lowest level. To visualize nucleoli, DsRed-B23 was co-transfected with EBNA-1 as a nucleolar marker. EBNA-1 nucleolar staining was detected in 78.6% of control siRNA–transfected HeLa cells, compared with only 10.7% in siEBP2-transfected cells (Fig. 6C). This loss-of-function experiment reinforces the above observations and strongly supports the requirement of EBP2 for EBNA-1 nucleolar targeting.

### Effect of EBNA-1 on the nucleolar transcription and protein synthesis

A wide range of viruses are known to target the nucleolus as a strategy to interfere with host-cell functions. The primary role of the nucleolus is the synthesis of rRNAs, which is carried out by RNA polymerase I–mediated transcription of ribosomal genes (rDNAs) (46). To determine whether nucleolar localization of EBNA-1 influences this process, we analyzed rRNA synthesis in situ by monitoring rRNA transcription. This was achieved through incorporation of BrUTP into nascent transcripts, followed by indirect immunofluorescence as previously described (47). The EBNA-1-NoLS* mutant, deficient in nucleolar localization, was used as a reference loss-of-function control to investigate the effect of EBNA-1 nucleolar targeting. The markedly reduced BrUTP fluorescence staining (Fig. 7A, panel a), compared to that observed with EBNA1-NoLS* (Fig. 7A, panel e), suggests a suppressive effect of EBNA-1 nucleolar targeting on rRNA synthesis. To validate this effect, ∼100 nuclei per condition were analyzed by 3D imaging using Tools for Analysis of Nuclear Genome Organization (TANGO) plugin in ImageJ (48), as shown in Fig. 7B (panels a–d). Quantification of nucleolar number per nucleus, mean fluorescence intensity, and volume enabled calculation of the fluorescence integrated intensity, which reflects total rRNA synthesis per cell. As shown in Fig. 7B (panel e), EBNA-1 nucleolar localization resulted in a 54% reduction in rRNA synthesis compared to the EBNA-1-NoLS* reference control.

**Fig. 7.**
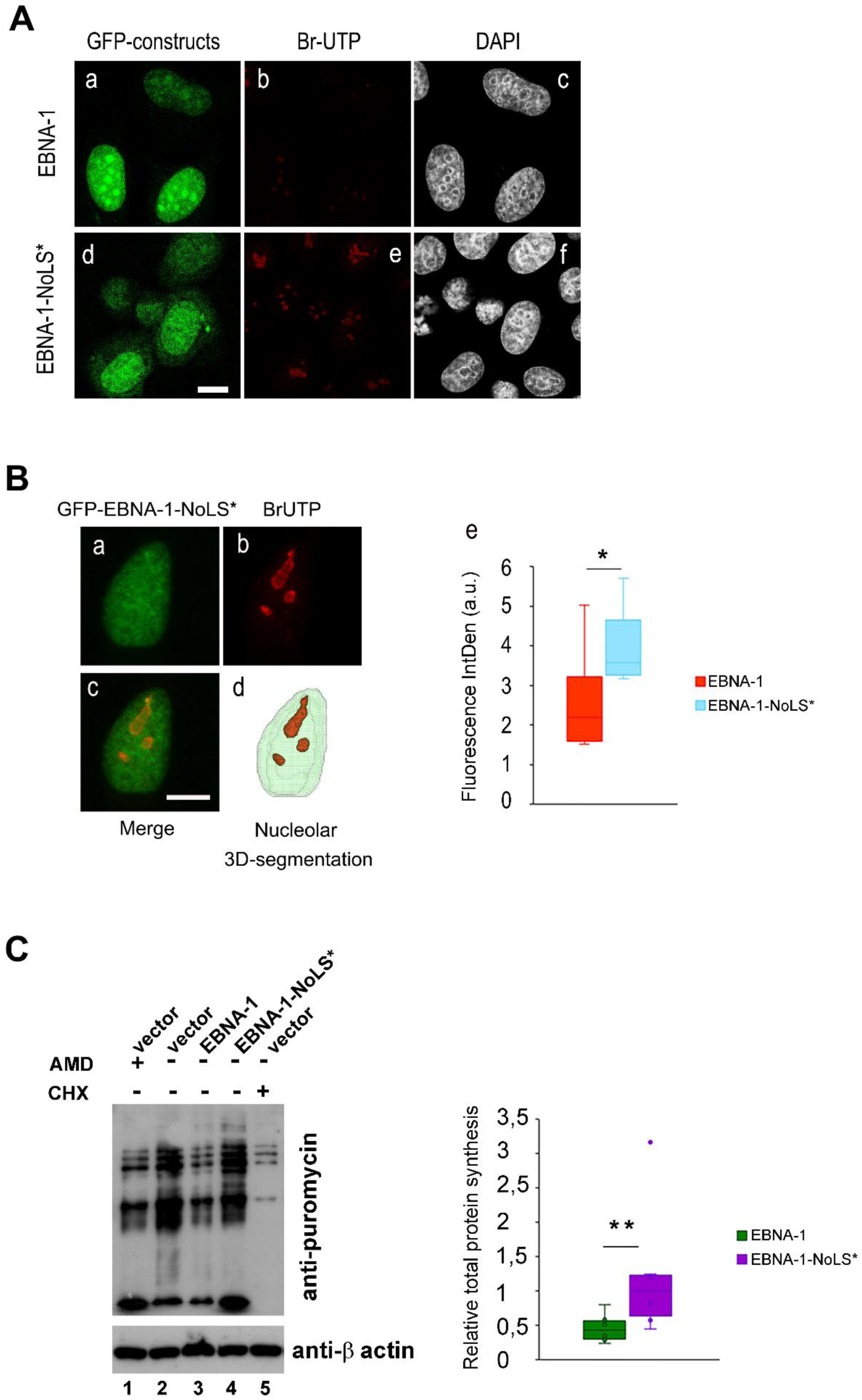
EBNA-1 nucleolar localization reduces rRNA and protein synthesis. (A) Hela cells expressing EBNA-1 or EBNA-1 NoLS* were cultured in medium containing Br-UTP for 5 min and processed to anti-BrdU indirect immunofluorescence to reveal rRNA synthesis by confocal microscopy observation. Single confocal z-sections are shown. Scale bars, 10 μm. (B) Schematic of the 3D nuclear and nucleolar imaging analysis by using TANGO pluging in ImageJ (*48*). First, nuclei are segmented from GFP-EBNA-1 staining (panel a), then BrdU staining within the subnuclear structures (panel b) can be segmented, visualized (panel d) and analyzed for spatial statistics, as represented by nucleolar fluorescence intensity histogram (panel e). Scale bars 10 μm. For each condition, 100 cells have been analyzed from 6 independent experiments. Data are presented as median (min–max). Comparisons between two groups were performed using Student’s *t*-test. *p-value < 0.05. (C) Hela cells expressing EBNA-1 or EBNA-1 NoLS*, treated with AMD or CHX or not treated, were cultured in medium containing puromycin for 10 min, harvested and cell lysates processed to anti-puromycin Western blot. Densitometric analyses from three independent experiments are shown as histograms. Data are presented as median (min–max). Comparisons between two groups were performed using Student’s *t*-test. **p-value < 0.01.

To assess the impact of this inhibition on ribosome assembly, we quantified protein synthesis as an indirect measure of ribosomal function. To this end, cells were exposed to a 10-min puromycin pulse. Puromycin, a nucleoside antibiotic that mimics aminoacyl-tRNA, causes premature termination of translation and remains covalently attached to the C terminus of nascent polypeptides, which can then be detected by anti-puromycin Western blotting (Fig. 7C). EBNA-1 expression led to an inhibition of protein synthesis comparable to that induced by AMD, an RNA polymerase I inhibitor, but less pronounced than that observed with CHX a protein synthesis inhibitor. In contrast, protein synthesis was not affected in cells transfected with EBNA-1-NoLS*, which is unable to enter nucleoli. These results indicate that EBNA-1–mediated inhibition of rRNA synthesis upon nucleolar entry reduces ribosome production and, indirectly, decreases cellular protein synthesis by ∼50%, as quantified by Western-blot densitometry and β-actin normalized.

### EBNA-1 nucleolar localization triggers oxidative stress–dependent inhibition of rRNA synthesis

Nucleoli also function as stress sensors, responding to DNA damage, heat shock, or viral infection (34). Such stresses increase reactive oxygen species (ROS), mainly H₂O₂, which diffuse into the nucleolus and inhibit RNA polymerase I activity (46). To test whether nucleolar targeting of EBNA-1 triggers oxidative stress, cells transfected with EBNA-1 or EBNA-1-NoLS* were co-transfected with HyPer7-NLS, an ultrasensitive H₂O₂ probe. This probe, derived from a circularly permuted YFP fused to an NLS and integrated into the OxyR domain of *Neisseria meningitidis* (49), shows increased green fluorescence under oxidative conditions when excited at 488 nm. Confocal microscopy revealed that HyPer7-NLS localized to both nuclei and nucleoli (Fig. 7A, d-f). In cells where EBNA-1 localizes to the nucleoli (Fig. 8A, a and c, arrow heads), Hyper7 staining was markedly stronger in both compartments compared with EBNA-1-negative cells (outlined with dashed lines) or with cells expressing EBNA-1-NoLS (Fig. 8A, b and d) excluded from the nucleoli. Treatment with the antioxidant N-acetyl-cysteine (NAC) reduced nuclear and nucleolar ROS staining to basal levels (data not shown) and restored rRNA synthesis, as measured by BrUTP incorporation (Fig. 8B), thereby demonstrating that ROS mediate the EBNA-1–induced decrease in rRNA synthesis upon nucleolar entry.

**Fig. 8.**
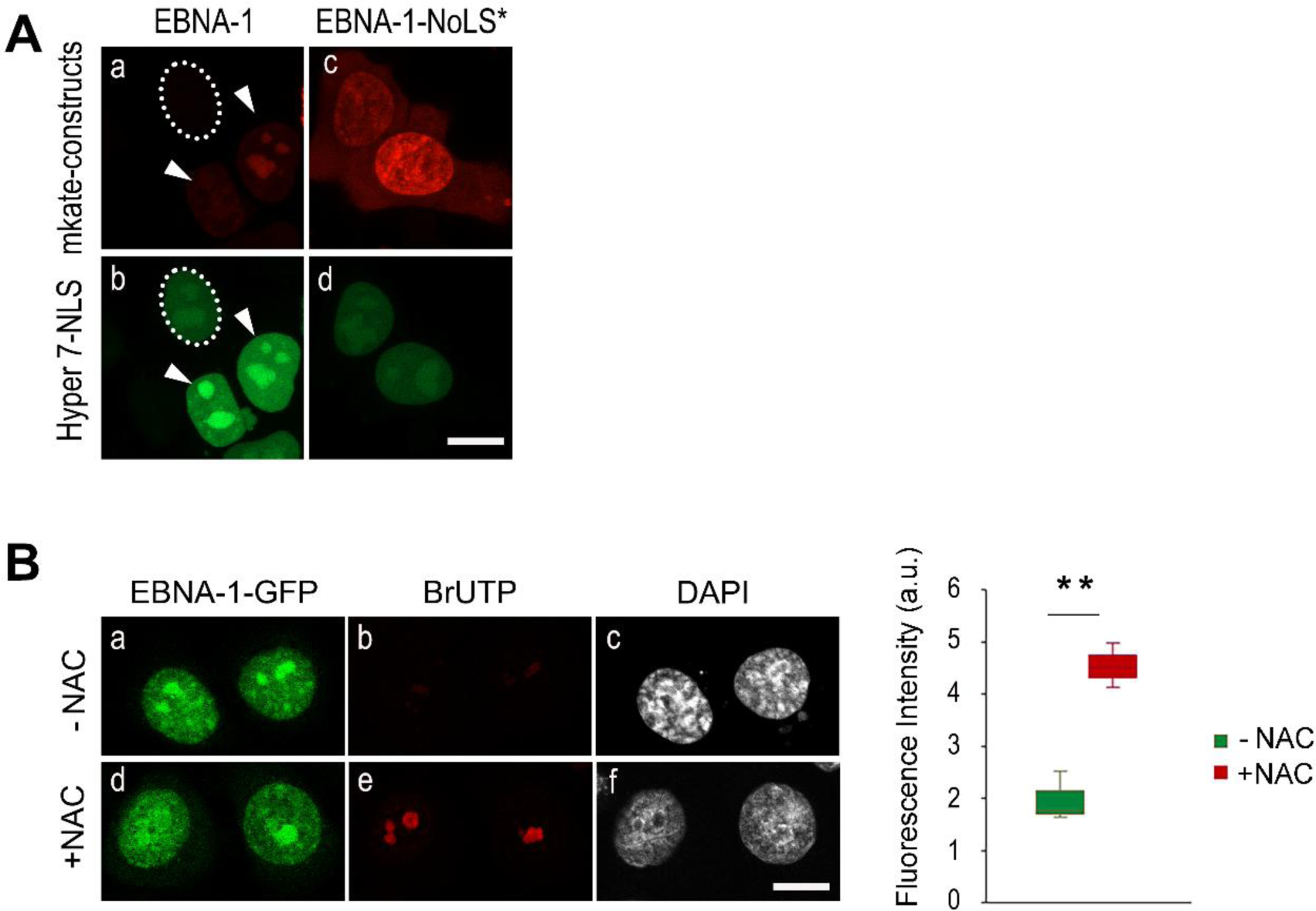
EBNA-1 nucleolar localization triggers oxidative stress at the origin of rRNA synthesis inhibition. **(A)** HeLa cells co-transfected with EBNA-1 or EBNA-1-NoLS* and Hyper 7-NLS were observed 24 h post-transfection confocal microscopy. Cells not expressing EBNA-1 (outlined with a dashed line), expressing EBNA-1 in nucleoli (arrowheads) or expressing EBNA1-NoLS* were compared for differences in Hyper7-NLS fluorescence intensity. Images show single confocal z-section. Scale bar, 10 μm. **(B)** HeLa cells expressing EBNA1-GFP were left untreated (−NAC) or treated with NAC for 20 h (+NAC), incubated in Br-UTP–containing medium for 5 min, and processed for indirect immunofluorescence using anti-BrdU antibody to visualize rRNA synthesis by confocal microscopy (Single z-sections are shown. Scale bars, 10 μm). For each condition (−NAC and +NAC), 50 images from three independent experiments were analyzed for 3D nuclear and nucleolar imaging using TANGO in ImageJ. BrdU fluorescence intensities are shown as histograms. Data are presented as median (min–max). Statistical comparisons between groups were performed using Student’s *t* test (**p < 0.01).

## DISCUSSION

Our study elucidates the mechanism underlying EBNA-1 nucleolar localization. This process is cell cycle–dependent and requires a bipartite nucleolar localization signal (NoLS) composed of tandem Weber motifs, as well as an interaction with the nucleolar protein EBP2. Nucleolar localization of EBNA1 leads to a ROS-dependent inhibition of rRNA synthesis, resulting in a global reduction of cellular protein production.

The nucleolus is a membrane-less organelle formed through liquid–liquid phase separation of its components from the surrounding nucleoplasm (50). Consequently, it lacks a dedicated active transport system comparable to the nuclear pore complex. Instead, nucleolar localization of proteins depends on interactions with nucleolar constituents such as rRNA or resident proteins including nucleolin, B23, or fibrillarin. Consequently, NoLSs do not conform to a recognized canonical sequence motif, but instead depend on a charged interaction mechanism that varies depending on the specific nucleolar molecule involved in the interaction.(51, 52). Notably, Arg/Gly-rich or Arg/Lys-rich regions are frequently involved in nucleolar targeting, as their high density of basic residues positively charged facilitates electrostatic interactions with negatively charged rRNA or acidic nucleolar proteins (52).

For instance, the Arg/Gly-rich region (GGRRGRRRGRGRGG) present in the HSV-1 ICP27 protein functions as a NoLS (53) through an RNA-binding mechanism (20). Similarly, Arg/Gly-rich regions can act as NoLSs via interactions with nucleolar proteins, as observed for the NoLS of the Japanese encephalitis virus core protein (DGRGPV) and the groundnut rosette virus ORF3 protein (RPRRRAGRSGGMDPR), which bind to B23 and fibrillarin, respectively. The GAR domain of nucleolin is essential for its efficient localization in the nucleolus (43, 54–56) and has been reported to interact with various host factors, including single- and double-strand RNA and DNA (42) and ribosomal proteins (41). Likewise, fibrillarin binds directly and specifically to small nucleolar RNA (57) and to Nop 56 (58). Although its GAR domain directly binds to RNA helicase p68 in nascent nucleoli during late telophase (59), it is dispensable for a correct nucleolar targeting of fibrillarin during interphase (58). These two examples indicate that GAR domain may play different role in nucleolar localization depending on the protein and/or of different stages of the nucleoli assembly.

EBNA1 has been shown to bind both cellular and viral RNAs *in vitro* through three independent RGG motifs (60), located between amino acids 33–56, 330–350, and 354–377, respectively (60). Our bioinformatic analysis revealed strong sequence similarity between the 330–350 aa region of EBNA-1 and the GAR domains of nucleolin and fibrillarin (see Fig. 3). However, deletion of this region did not impair EBNA-1 nucleolar localization, clearly indicating that it does not function as a NoLS. Similarly, the 33–56 RGG motif does not fulfill this role, as several nucleolar-excluded mutants (see Fig. 4) still retain this region. Regarding the third RGG motif (aa 354–377), our data indicate that it only partially overlaps the EBNA-1 NoLS, since its deletion still allows nucleolar localization of EBNA1 in approximately 17% of transfected cells (see Fig. 4). Taken together, these results indicate that EBNA-1 binding to RNA via RGG-enriched motifs is not involved in its nucleolar targeting mechanism.

Another class of basic amino acid motif, enriched in lysine residues and organized around an R/K–R/K–X–R/K pattern, has also been implicated in the nucleolar targeting of proteins. First identified by Weber *et al*. as the NoLS of cellular proteins ARF and Hdm2 (44), it was subsequently found in the nucleolar protein Nop 25 (61) and in several ribosomal proteins, including S25 (62), L7a (63), L22 (35) and S6 (64). Notably, Weber sequences are found in many viral proteins that associate with the nucleolus. In some cases, a single Weber sequence is sufficient for nuclear localization. Examples include the regulator protein Rex from HTLV1 (65), both the large and small antigens from hepatitis delta virus (HDV) (66), as well as Nucleoproteins of influenza A virus (67), salmon anemia virus (68), SARS-CoV-1 (69) and porcine reproductive and respiratory syndrome virus (PRRSV) (70). In other cases, multiple adjacent or overlapping Weber motifs form a functional cluster, as seen in HSV-1 γ₁34.5 protein (71), African Swine Fever Virus (SFV) I14L protein (72), bovine herpesvirus-1 BICP27 protein (73), HIV1 Tat and Rev (74, 75) and feline coronavirus 3b protein (76). Two distinct Weber motifs that independently drive nucleolar localization have also been described in the Semliki forest virus capsid and adenovirus protein V (77, 78). Conversely, two Weber sequences, separated by a linker but acting cooperatively to mediate nucleolar targeting, have been identified in the CHIKV capsid (79), the african swine fever virus I14L protein (72), the Marek disease virus oncoprotein MEQ (80) and the potato virus A VPg domain of NIa protein (81). Consistent with these observations, we identified a similar bipartite NoLS in the EBNA-1 sequence. Interestingly, this NoLS overlaps with the previously characterized NLS of EBNA-1 (82), a configuration repeatedly reported for nucleolar proteins (reviewed in reference (83). Moreover, by integrating experimentally validated NoLS datasets with NLS prediction tools, Kurnaeva *et al.* (84) further demonstrated that such overlaps occur more frequently in viral than in human proteins. Despite this partial overlap, the mutation affecting both the NoLS and the NLS does not sufficiently impair NLS function to prevent EBNA-1 nuclear import. However, it may account for the cytoplasmic staining observed with the EBNA-1-NoLS* mutant, a feature not detected in any of the other mutants we generated.

The nucleolar staining pattern of EBNA1, which uniformly spans the entire nucleolus (Fig.1, panel 1), is characteristic of GC localization, where late rRNA processing occurs through proteins such as B23 and EBP2. This pattern therefore supports an interaction between EBNA1 and EBP2. With the exceptions of HSV1-ICP27 or HIV-Tat which has been shown to bind RNA (20, 51, 52), the nucleolar components identified so far as viral protein partners include nucleolin (28, 66, 68, 84–89), B23 (90–98) and fibrillarin (81, 99, 100). Extending these observations, we found that mutations abolishing EBNA-1 nucleolar localization also disrupt its interaction with EBP2, demonstrating that EBP2 binding is critical for nucleolar targeting. To our knowledge, this is the first report implicating EBP2 in the nucleolar targeting of a viral protein.

Using live-cell imaging approaches, a previous study conducted in our laboratory challenged the proposed chromosomal tethering role attributed to the EBNA-1/EBP2 interaction (15). In the present work, the application of the same imaging strategies allowed us to further define the biological significance of this interaction and to establish its relationship with EBNA-1 nucleolar localization, the subsequent induction of oxidative stress, and the inhibition of rRNA synthesis. One might argue that physiological relevance of experiments based on overexpression of fluorophore tagged EBNA-1 and EBP2 may therefore be questioned. Indeed, whether EBNA-1 displays a comparable nucleolar localization during natural EBV latency in infected B lymphocytes or epithelial cells, and what functional consequences such localization might entail, remain to be determined. However, use of fluorophore-tagged EBNA-1 in transfected cells provides experimental advantages that are not readily achievable in the context of viral infection. In particular, the precise subnuclear distribution of EBNA-1 cannot be reliably assessed using immunofluorescence, as chemical fixation is known to alter the localization of nuclear proteins. A transfection-based system also allows the specific contribution of EBNA-1 to be examined independently of other viral proteins or noncoding RNAs expressed during EBV latency. In addition, the generation and functional analysis of EBNA-1 mutants would be technically challenging within the full viral genome and could raise biosafety concerns if certain mutations were to enhance viral pathogenic potential. Accordingly, although this system does not fully recapitulate the physiological context of EBV latency, it currently provides the most suitable experimental framework to investigate EBNA-1 subnuclear dynamics and to dissect its specific functional contributions under controlled conditions.

Our data demonstrate that EBNA1 nucleolar localization is tightly regulated in a cell cycle–dependent manner. Among viral proteins, such regulation has previously been described for the nucleocapsid (N) protein of avian coronavirus IBV, which accumulates in the nucleolus during G2/M (101). In contrast, EBNA-1 localizes to the nucleolus mainly during S phase. Given that rRNA transcription peaks during S and G2 phases [reviewed in (46)], a virus-induced perturbation of nucleolar function during S phase could profoundly impact ribosome biogenesis and, consequently, cellular physiology, including growth, metabolism, and cell cycle progression. rRNA synthesis, the first step of ribosome biogenesis, is markedly influenced by EBNA-1 and, more specifically, by its nucleolar localization (Fig. 7 A and B). Indeed, cells expressing the EBNA1-NoLS* mutant, which retains chromatin association but fails to enter the nucleolus, exhibit rRNA synthesis to levels comparable to those of GFP-transfected controls (data not shown).

EBNA-1 is the only viral protein consistently detected in all EBV-associated tumors, including Burkitt lymphoma and nasopharyngeal carcinoma (NPC), suggesting a key role in the virus’s oncogenic and transforming potential. Inhibition of rRNA and protein synthesis cannot account for the oncogenic properties of EBNA-1, since cancer cells, on the contrary, require increased rRNA and protein synthesis to sustain proliferation. For example, the SV40 T antigen, a well-known oncogenic protein, enhances rRNA synthesis (102). However, given the deleterious effects of ROS on genome stability, the oxidative stress triggered by EBNA-1 nucleolar localization, which leads to rRNA synthesis inhibition, may contribute to the oncogenic properties of EBNA-1. Indeed, Gruhne *et al.* reported that EBNA-1 triggers chromosomal aberrations, DNA double-strand breaks, and activation of the DNA damage response (DDR) in B lymphocytes. These manifestations of genomic instability are ROS-dependent and can be reversed by antioxidant treatment (103). Long term EBNA-1 expression in NPC have also been reported to increase ROS production and upregulate the NADPH oxidase NOX1 and NOX2, known to generate ROS (104). The present study provides new insights into the transforming mechanism of EBNA-1 by showing that EBNA1, but not the EBNA1-NoLS* mutant, increases ROS levels within both the nucleus and the nucleolus. These findings indicate that the transforming potential of EBNA-1, mediated through oxidative stress induction, depends on its nucleolar localization.

Nucleolar stress signaling is characterized by rRNA synthesis inhibition, delocalization of ribosomal proteins (RP) to the nucleoplasm, RP binding to MDM2 leading to P53 activation and subsequent cell cycle arrest, senescence or apoptosis (105). Several viruses have been reported to interact with nucleolar stress signalling. SARS-CoV nucleoprotein localizes to the nucleoli of infected cells (106) and interacts with B23 (96), inducing inhibition of B23 phosphorylation and subsequent cell-cycle arrest (96). In the case of influenza A virus, the non-structural NS1 protein localizes in nucleoli and inhibits rRNA synthesis (107). The West Nile virus capsid protein induces p53-mediated apoptosis via the sequestration of HDM2 into the nucleolus (108). Finally, the Zika virus capsid protein (ZIKV-C) has been reported to localize in the nucleoli, causing nucleolar disruption marked by the translocation of B23 to the nucleoplasm, which leads to the activation of P53 and induction of apoptosis in neural cells (109).

Paradoxically, despite inhibiting rRNA synthesis, EBNA-1 is known to confer resistance to apoptosis. This suggests that EBNA-1 may interfere with the nucleolar stress signaling pathway downstream of rRNA synthesis inhibition. This hypothesis aligns with findings from Frappier and colleagues, showing that EBNA-1, through its interaction with the deubiquitinating enzyme USP7/HAUSP, disrupts USP7 binding to p53, thereby promoting p53 proteasomal degradation (12, 110) and the disassembly of PML nuclear bodies (111), both of which contribute to apoptosis resistance. Additionally, survivin which is negatively regulated by P53, is positively regulated by EBNA-1, in EBV-associated B-lymphoma cells, by an indirect binding at the surviving promotor (13). Through its effects on p53 and survivin, EBNA-1 promotes host cell survival and thereby may ensure EBV episome maintenance. Yet, these same activities impair repair, senescence, and apoptotic pathways, allowing oxidative stress–induced genome instability to persist and thereby creating a context conducive to oncogenesis.

Altogether, our results establish nucleolar targeting of EBNA1 as a multifaceted process, mediated by both NoLS motifs and EBP2 binding, and linked to ROS-dependent nucleolar stress. Future work should determine whether the “dissociation” between nucleolar stress and p53 activation contributes to EBNA-1-driven oncogenesis.

## MATERIALS AND METHODS

### Recombinant Plasmids

The DsRed-B23 construct was reported previously (58). Recombinants plasmids GFP-EBP2 and DsRed-EBP2 that express EBP2 inserted in pEGFP-C1 and pDsRed1-C1 (BD Bioscience Clontech), respectively, as well as EBNA-1-GFP and EBNA-1-DsRed, that express EBNA-1 construct deleted from the central GlyAla repeated region (aa 93 to 325) and inserted in pEGFP-N1 or pDsRed-N1, respectively, were described earlier (15).

The EBNA-1 Δ325-349 or Δ350-376 or Δ325-376 deletion constructs were obtained by using the EBNA-1-GFP plasmid from which each aa-indicated region was removed by using a two-step PCR overlapping procedure. In short, the sequence to be deleted is called part B, the upstream sequence part A and the downstream sequence part C. All primers are described in table S1-A. In a first PCR step, primers 1-Forward (PCR1-F) and 1-Reversed (PCR1-R) were used to amplify the part A whereas primers PCR2-F and PCR2-R were used for amplify the part C of EBNA-1. Primer PCR1-R was drawn to overlap part C and primer PCR2-F to overlap part A. The partly overlapping PCR products from the two separate reactions were gel purified and mixed together with primers PCR1-F and PCR2-R in a second PCR in order to produce DNA encoding the part A fused to part C (A-C) of EBNA1. This PCR product from the second reaction (A-C sequence) was digested with EcoR1 and Apa1 and cloned into EBNA-1-GFP plasmid in place of A-B-C sequence.

Site-directed mutagenesis was performed to substitute RRGR or KRPR EBNA-1 weber’s motifs in AAGR or AAPR respectively, by using EBNA-1-GFP plasmid as a template and a two-step PCR overlapping procedure with primers listed in table S1-B. Briefly, point mutations were introduced by reversed primer of the first PCR (PCR1-R) that partially overlap with the forward primer of the second PCR (PCR2-F). The partly overlapping PCR products from the two separate reactions were gel purified and mixed together with primers PCR1-F and PCR2-R in a second PCR in order to produce DNA encoding AAGR or AAPR point mutation in EBNA1 sequence. These PCR product from the second reaction (containing AAGR or AAPR) were digested with EcoR1 and Apa1 and cloned into EBNA-1-GFP plasmid in place of non-mutated sequence.

The EBNA-1 AAGR/AAPR double mutant was constructed on the same principle by substituting the KRPR motif for AAPR via site-directed mutagenesis of the EBNA-1-AAGR-GFP plasmid.

The EBNA-1-mKate and EBNA-1-NoLS*-mKate constructs were obtained by fusing the EBNA-1 derivatives to mKate2 far-red fluorescent protein (112). The EBNA-1-GFP constructs were digested by BamHI and NotI and replaced by the corresponding mKate2 sequence from the pmKate2-N vector (a gift from Dmitry Chudakov).

### Cell culture, DNA transfection and cell cycle synchronization

Human Hela cells (ATCC:CCL2) were cultured in Dulbecco’s modified Eagle’s medium (DMEM) supplemented with 10% foetal calf serum (FCS) and 2mM L-Glutamine in a 95% air-5% CO-2 incubator at 37°C. For DNA transfection experiments, Hela cells were seeded at a density of 150,000 cells on glass bottom μ-dish (Ibidi) until they reached approximately 50% confluence. Plasmids were transfected with Effectene Transfection Reagent (Qiagen) according to manufacturer’s recommendations. Cells were observed by confocal microscopy 24 h post-transfection. For prenucleolar bodies (PNBs) observation, HeLa transfected cells were blocked in prometaphase by nocodazole (0.04 µg/ml for 4 h), selectively harvested by mechanical shock, washed and resuspended in nocodazole-free medium containing AMD (0,1 μg/ml) to specifically inhibit rRNA transcription during 2 h and then were observed by confocal microscopy. For cell cycle synchronization at G1/S border, a double block with 2′-deoxythymidine (dT, Merck) has been performed as previously described (113). Briefly, transfected Hela cells were incubated in DMEM containing 3 mM dT for 15 h, washed 3 times and incubated in a fresh medium for 9h and again for 15h in a medium with 3 mM dT. Cells were washed, incubated in a fresh medium and observed by confocal microscopy immediately for late G1 phase, after 3 h incubation for S phase and after 6 h incubation for G2 phase.

### Assay of rRNA transcription in situ

The assay was performed by labelling neosynthetised rRNAs by 5-Bromo-2′-uridine 5′-triphosphate (BrUTP) incorporation in transfected HeLa cells grown on glass coverslips. BrUTP was introduced into cells by a hypotonic shock. Briefly, 24h post transfection, cells where washed with a hypotonic buffer (30 mM KCl, 10 mM Hepes) and a drop of this buffer containing 10 mM BrUTP was applied upon each coverslip for 10 min incubation. Cells were then washed and incubated on a normal medium for 30 min. After cell fixation with methanol for 20 min at −20°C, air-drying for 5 min, BrUTP incorporation was detected as previously described (114) by immunofluorescence labeling using mouse anti-BrdU antibodies (Merck) revealed by the Alexa-Fluor-594-conjugated anti-mouse antibodies. Coverslips were mounted in Fluoroshield with DAPI (4’, 6’-diamino-2-phenylindole) and viewed for confocal analysis.

### Confocal microscopy and cell imaging

Confocal analyses were performed using an inverted Leica SP5 microscope equipped with a thermostatic chamber (37°C, 5% CO₂) and a 63× oil immersion Plan-Apochromat objective (NA 1.4). Cells were scanned at 400 Hz with a resolution of 1024x1024 pixels and a voxel size of 107 nm x 107 nm x 300 nm. The pinhole was adjusted so as to fit the airy disc. DAPI, GFP and DsRed were respectively excited by a 405 nm 488 nm and 561 nm wavelength, and emission fluorescences were detected respectively by photomultiplicator (PM) between 410-480 nm, 495-550 nm, 590-670 nm. Three channels were recorded sequentially at each Z-step. Z series projections, merge images and 3D visualizations were performed using ImageJ software [W. S. Rasband ImageJ (U.S. National Institutes of Health, Bethesda, MD; http://rsb.info.nih.gov/ij, 1997-2009)]. Nuclear and nucleolar 3D quantitative analyses were performed using Tools for Analysis of Nuclear Genome Organization (TANGO) (48) and visualized with the 3D Viewer plugin in ImageJ.

FRET analyses between GFP-EBP2 and either DsRed-EBNA-1 or DsRed-EBNA-1-NoLS were performed by exciting GFP at 488 nm, and fluorescence emission spectra (488–648 nm) were collected using a Leica HyD GaAsP hybrid detector (Leica Microsystems).

### Protein synthesis assay

Protein synthesis was estimated by using a nonradioactive method in which puromycin is incorporated into nascent proteins and causes termination (115, 116). Briefly, 48 h post-transfection, cells were pulse-labelled with 9 μM puromycin and incubated for 10 min at 37°C in a 5% CO₂ atmosphere. Control cells were treated with 8 nM Actinomycin D (AMD) or 50 μM cycloheximide (CHX), 3 h or 10 min before pulse labeling, respectively. The cell lysates were prepared with RIPA lysis buffer, total protein concentrations were determined using the Pierce BCA Protein Assay Kit (Thermo Scientific) and 10 μg of the total amount of pulse-labelled proteins were separated by SDS-PAGE (6% acrylamide/bisacrylamide 37,5:1) and electrically transferred onto a PVDF membrane (Hybond-P, Amersham). The membrane was blocked in PBS-T buffer (0,1% Tween 20) containing 5% nonfat dry milk at room temperature for 1 h followed by incubation with the primary antibodies (dilution 1:10,000 with 1% BSA for anti-puromycin (mAb, Merck), and dilution 1/10,000 with 1% BSA for an anti-β−actin (C4 monoclonal Ab, Santa Cruz)) at room temperature during 1 h. β-actin was used to confirm equal protein loading.

After three washes with PBS-T buffer, the membrane was incubated at room temperature for 1 h with anti-mouse HRP-labelled secondary antibodies (Cytiva, 1:10,000 dilution) for both the anti-puromycin and anti-β-actin antibodies, and then revealed with an HRP substrate (SuperSignal West Pico PLUS, ThermoFisher). Protein synthesis was estimated based on the intensity of immunoreactive bands using ImageJ.

### Intracellular detection of Reactive Oxygen Species (ROS)

Hydrogen peroxide was monitored using HyPer7-NLS (Addgene #136468), an ultrasensitive H₂O₂ probe derived from a circularly permuted YFP fused to SV40-LT-NLS and integrated into the OxyR domain of *Neisseria meningitidis* (49). Cells were co-transfected with HyPer7-NLS and either EBNA-1-mKate or EBNA-1-NoLS-mKate. 24 h post-transfection, cells were analyzed by confocal microscopy. HyPer7-NLS was excited at 488 nm, and fluorescence was detected between 490 and 561 nm.

### Bioinformatic analyses

Nucleolar sequences were fetched from main databases based upon their IPI numbers (117) which are given in the Nucleolar Proteome Database NoPdb 3.0 (37, 118). The human proteome was extracted from the ENSEMBL system (119) (February 2009 Homo sapiens high coverage assembly GRCh37). Computations for human proteome without nucleolar proteins were achieved on a human proteome database without sequences having more than 95% identity with a nucleolar protein. Tripeptide motifs and hexapeptide motifs were searched by using the program fuzzpro from the EMBOSS (120) software suite (under the GNU Public License). The GR rich motifs were extracted and counted using a home-made program. A motif must have a minimum length of 7 amino-acids, it must be covered at 80% by G or R, and it must contain a minimum of 20% of G or R. Moreover, it must begin and end with GR or RG dipeptides.

### siRNA-mediated knockdown experiments

Small interfering RNAs (siRNAs) targeting human EBP2 (si-EBP2) were designed and synthesized by Horizon. Four distinct siRNA sequences (siRNA D-019834-01, siRNA D-019834-02, siRNA D-019834-03 and siRNA D-019834-04) were employed to ensure specificity and efficiency. A non-targeting scrambled siRNA (si-control) was used as a negative control. Hela cells were seeded in Ibidi or 6-well plates and transfected at approximately 30% confluence using Lipofectamine™ RNAiMAX Transfection Reagent (Invitrogen) according to the manufacturer’s instructions. Briefly, siRNAs (final concentration: 60 nM) were diluted in Opti-MEM™ (Gibco) and incubated with Lipofectamine™ RNAiMAX for 10 min at room temperature before adding to the cells. After 4 h of incubation, the medium was replaced with fresh complete culture medium, and cells were further incubated for 24-48 h. Knockdown efficiency of si-EBP2 was assessed at mRNA level by qRT-PCR (see qRT-PCR section) and at protein level by western blotting.

EBP2 Western blot analysis was performed by using a rabbit monoclonal antibody against human EBP2 (Thermo Fisher), a mouse monoclonal antibody against α-tubulin (Santa Cruz Biotechnology), and HRP-conjugated secondary antibodies (Cytiva) against rabbit or mouse IgG. Cellular protein extracts were prepared with RIPA lysis buffer and concentrations were determined using the Pierce BCA Protein Assay Kit (Thermo Scientific). 10 μg proteins were separated by SDS-PAGE (10% acrylamide/bis-acrylamide 37,5:1) and electrically transferred onto a nitrocellulose membrane (Amersham). The membrane was blocked in PBS-T buffer (0.1% Tween-20) containing 5% BSA at room temperature for 1 h followed by incubation with the primary antibodies (1:2000 dilution for EBP2 antibody and 1:20,000 for a-tubulin antibody) at 4°C overnight. After three washes with PBS-T buffer, the membrane was incubated at room temperature for 1 h with HRP-labelled secondary antibody (anti-rabbit 1:10,000 dilution). Detection by enzyme-linked chemiluminescence was performed according to the manufacturer’s protocol (Cytiva). Subsequently, three washes of the membrane with PBS-T buffer were done and the specific protein band were detected by enhanced chemiluminescence using the ChemiDoc Imager (BIO-RAD). Density of protein bands were quantified using Image Lab 6.1 (BIO-RAD).

### Quantitative RT-PCR Analysis

24 h, 48h and 72 h after si-RNA transfection, quantitative RT-PCR (qRT-PCR) analyses were performed as previously described (121). Briefly, total RNA was extracted from HeLa cells using a Total RNA Miniprep Kit (New England Biolabs, Évry, France), according to the manufacturer’s instructions. cDNA was synthesized from 1 µg total RNA using a ProtoScript^®^ II First Strand cDNA Synthesis Kit (New England Biolabs, Évry, France). Quantitative PCR was performed using Luna qPCR master mix (New England Biolabs, Evry, France). The specific primers are listed in supplementary table S2. Mastercycler^®^ RealPlex2 (Eppendorf, Évry-Courcouronnes, France) was used to perform amplification with the following thermal cycling conditions: denaturation at 95°C for 1 min followed by 40 cycles of denaturation at 95°C for 15 s, and annealing and elongation at 60°C for 45 s. A dissociation curve for each well was performed by running the following program: 95°C for 15 s, 60°C for 15 s and 60 to 95°C at 2°C/min. The obtained Ct (cycle threshold) values of the target genes were normalized to the NAPDH housekeeping gene, and the 2^−ΔΔCT^ method was used to calculate fold changes (122). Three biological replicates were performed for each gene (*n* = 3).

### Antioxidant treatment

Cells were preincubated for 20 h with 1 mM N-acetyl-L-cysteine (NAC; Sigma-Aldrich). NAC was removed during BrUTP incorporation, after which cells were further incubated for 30 min at 37°C and 5% CO₂ in DMEM supplemented with 1 mM NAC.

### Statistical analysis

For nucleolar 3D quantitative analysis and puromycin immunoblotting quantification, data are presented as median (min–max). Comparisons between two groups were performed using Student’s *t*-test. For EBP2 RT-qPCR and immunoblot quantification, data are presented as mean ± SD and statistically evaluated by one-way ANOVA followed by Tukey’s post hoc test, using R (R Foundation for Statistical Computing, Vienna, Austria). *p*<0.05; *p<0.01*.

## DATA AVAILABILITY

The data that support the findings of this study are available from the corresponding author upon request.

## ACKNOWLEDGMENTS

We thank the “Imagerie Paris Seine” imaging platform for confocal microscopy and FRET analyses. We thank Dmitriy Chudakov for pmKate2-N vector and Olivier Albagli for EBNA-1-mkate constructions and careful reading of the manuscript.

This study was supported by grant from the *Agence Nationale de la Recherche (ANR), France*. M.-M.C. received support from a scholarship from the French Embassy (Service for Cooperation and Cultural Action, SCAC) and from the “Young Talents Sub-Saharan Africa for Women in Science” Award, funded by the L’Oréal Foundation and UNESCO.

## SUPPLEMENTAL MATERIAL

Tables S1 and S2

