## Supplemental Tables S1 and S2 for "Nucleolar Targeting and ROS-dependent inhibition of rRNA synthesis by Epstein-Barr Virus Nuclear Antigen 1"

### Supplemental Material

| | primer name | primer sequence |
|---|---|---|
| **A-EBNA-1 deletion mutants** | | |
| Δ325–349 | PCR1-F | 5'-GCTTC*GAATTC*CCACC**ATGAC-3'** |
| | PCR1-R | 5'-GGCCTCCACT**TGCTCCTGTTCCACCG-3'** |
| | PCR2-F | **5'-GAACAGGAGCA**AGTGGAGGCCGCC-3' |
| | PCR2-R | 5'-GCG*GGGCCC*TGCTCTATCGCTC-3' |
| Δ350-376 | PCR1-F | 5'-GCTTC*GAATTC*CCACC**ATGAC**-3' |
| | PCR1-R | 5'-GCCTCTTTTCTCC **ACCTCCTCGACCCC-3'** |
| | PCR2-F | **5'-GGTCGAGGAGGT**GGAGAAAAGAG-3' |
| | PCR2-R | 5'-GCG*GGGCCC*TGCTCTATCGCTC-3' |
| Δ325-376 | PCR1-F | 5'-GCTTC*GAATTC*CCACC**ATGAC**-3' |
| | PCR1-R | 5'-CTCTTTTCTCC **TGCTCCTGTTCCACC-3'** |
| | PCR2-F | **5'-GAACAGGAGCA**GGAGAAAAGAGGCCCA-3' |
| | PCR2-R | 5'-GCG*GGGCCC*TGCTCTATCGCTC-3' |
| **B-EBNA-1 point mutation mutants** | | |
| RRGR--AAGR | PCR1-F | 5'-GCTTC*GAATTC*CCACCATGAC-3' |
| | PCR1-R | **5'-CGTCCTCTACC**CgcGgcGCCTCC-3' |
| | PCR2-F | 5'-**GGTAGAGGA**CGTGAAAGAGCC-3' |
| | PCR2-R | 5'-GCG*GGGCCC*TGCTCTATCGCTC-3' |
| KRPR--AAPR | PCR1-F | 5'-GCTTC*GAATTC*CCACCATGAC-3' |
| | PCR1-R | **5'-GGACTCCTGGG**CgcCgcTTCTCC-3' |
| | PCR2-F | **5'-CCCAGGAGTCC**CAGTAGTCAGTCA-3' |
| | PCR2-R | 5'-GCG*GGGCCC*TGCTCTATCGCTC-3' |

**Table S1.** Forward (-F) and reversed (-R) primers used **A**-for EBNA-1 deletion strategies: Boldface letters indicate part A sequence, underlining indicates part C sequence, while italics indicate restriction sites (EcoR1 for primer PCR1-F, Apa1 for primer PCR2-R). **B-** for EBNA-1 site directed mutagenesis strategies: Boldface letters indicate overlapping sequences, mutated nucleotides are written in lower case characters, while italics indicate restriction sites (EcoR1 for primer PCR1-F, Apa1 for primer PCR2-R).

| Genes | Gene accession number | Forward Primer 5'-3' | Reverse Primer 5'-3' |
|---|---|---|---|
| **EBP2** | NM_006824.3 | AGAAAGGCTCAAAGTGGAACA | CATGACACCAACAGAAGGGAT |
| **GAPDH** | NM_002046.7 | TGCACCACCAACTGCTTAGC | GGCATGGACTGTGGTCATGAG |

**Table S2.** List of primers used for quantitative real-time PCR.
